# qqman: an R package for visualizing GWAS results using Q-Q and manhattan plots

**DOI:** 10.1101/005165

**Authors:** Stephen D. Turner

## Abstract

**Summary:** Genome-wide association studies (GWAS) have identified thousands of human trait-associated single nucleotide polymorphisms. Here, I describe a freely available R package for visualizing GWAS results using Q-Q and manhattan plots. The `qqman` package enables the flexible creation of manhattan plots, both genome-wide and for single chromosomes, with optional highlighting of SNPs of interest.

**Availability:** `qqman` is released under the GNU General Public License, and is freely available on the Comprehensive R Archive Network (http://cran.r-project.org/package=qqman). The source code is available on GitHub (https://github.com/stephenturner/qqman).

## 1 INTRODUCTION

Genome-wide association studies (GWAS) have been successful in identifying thousands of trait and disease-associated single nucleotide polymorphisms (SNPs) (Welter et al., 2014). A typical GWAS will examine hundreds of thousands to millions of polymorphic variants in many samples, testing their statistical association with discrete outcomes (a “case-control study,” such as diabetic vs. non-diabetic) and/or continuous outcomes (quantitative traits such as height, lipid levels, or expression level of a gene). The primary result of this statistical analysis is a list of SNPs, their associated chromosomal position, and a *P*-value representing the statistical significance of the association.

A commonly used method used to visualize GWAS results is the “manhattan plot” – a plot of the –*log*_10_(*P*-value) of the association statistic on the *y*-axis versus the chromosomal position of the SNP on the *x*-axis. Regions with many highly associated SNPs in linkage disequilibrium appear as “skyscrapers” along the plot. Another commonly used results diagnostic plot is the quantile-quantile (“Q-Q”) plot. Q-Q plots display the observed association *P*-value for all SNPs on the *y*-axis versus the expected uniform distribution of *P*-values under the null hypothesis of no association on the *x*-axis. Strongly associated SNPs will deviate from the diagonal at the upper-right end of the plot, while systematic deviation from the diagonal may indicate problems with the data, such as population stratification or cryptic relatedness.

One of the most commonly used software packages for manipulating and analyzing GWAS data is PLINK (Purcell et al., 2007). Here, I describe an R package that allows for quick and flexible generation of publication-ready Q-Q and manhattan plots directly from PLINK results files.

## 2 IMPLEMENTATION & EXAMPLE USAGE

The `qqman` software is developed as a package for the R statistical computing environment, and is released under the GNU General Public License on the Comprehensive R Archive Network (CRAN). The `qqman` package ships with an example set of GWAS results called `gwasResults` in a format similar to the output from `plink --assoc` that will be used in the examples below. For more examples, please view the package vignette.

The `manhattan()` function in the `qqman` package takes a data frame with columns containing the chromosome number, chromosomal position, *P*-value, and optionally the SNP name. By default, `manhattan()` looks for column names corresponding to those output by the `plink --assoc` command, namely, “CHR,” “BP,” “P,” and “SNP,” although different column names can be specified by the user. Calling the `manhattan()` function with a data frame of GWAS results as the single argument draws a basic manhattan plot, defaulting to a black-and-white color scheme:

~~~
# Load pkg, view data, draw basic manhattan plot
library(qqman)
head(gwasResults)
manhattan(gwasResults)
~~~

Further graphical parameters can be passed to the `manhattan()` function to control things like point character, size, colors, etc. When run with defaults, the `manhattan()` function draws horizontal lines drawn at –*log*_10_(1×10^-5^) for “suggestive” associations and –*log*_10_(5×10^-8^) for the “genome-wide significant” threshold. These can be moved to different locations or turned off completely with the `suggestiveline` and `genomewideline` arguments, respectively.

~~~
# Reduce point size to 80%
manhattan(gwasResults, cex=0.8)
# Change colors to alternating blue and orange
manhattan(gwasResults, col=c(“blue4”, “orange3”))
# Turn off the suggestive line at 1e-5:
manhattan(gwasResults, suggestiveline=FALSE)
~~~

The behavior of the `manhattan()` function changes slightly when results from only a single chromosome are used. Here, instead of plotting alternating colors and chromosome number on the x-axis, the SNP’s position on that chromosome is plotted on the x-axis:

~~~
# Plot only results on chromosome 3
chr3 <- subset(gwasResults, chr==3)
manhattan(chr3)
~~~

The `manhattan()` function has the ability to highlight SNPs of interest. Here, a character vector containing SNP names to highlight. The `qqman` package ships with an example set of “interesting” SNPs called `snpsOfInterest`, used in the example below.

~~~
# View and highlight SNPs of interest
head(snpsOfInterest)
manhattan(gwasResults, highlight=snpsOfInterest)
~~~

Additionally, highlighting SNPs of interest can be combined with limiting to a single chromosome to enable “zooming” in to a particular region containing SNPs of interest:

~~~
# View and highlight interesting SNPs on chr3:
chr3 <- subset(gwasResults, chr==3)
manhattan(chr3, highlight=snpsOfInterest)
~~~

Finally, the `qq()` function can be used to generate a Q-Q plot to visualize the distribution of association *P*-values. The `qq()` function takes a vector of *P*-values as its only required argument. Further graphical parameters can be passed to `qq()` to control the plot title, axis labels, point characters, colors, point/text sizes, etc.

~~~
# Draw a Q-Q plot with a title
qq(gwasResults$P, main="Q-Q plot of P-values”)
~~~

**Fig. 1.**
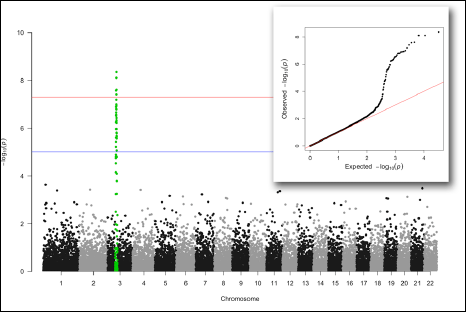
Manhattan plot highlighting SNPs of interest on chromosome 3, with Q-Q plot showing substantial deviation from the diagonal (inset).

## 3 CONCLUSIONS

The `qqman` package is a user-friendly tool to visualize results from GWAS experiments using Q-Q and manhattan plots. These graphics can be created in other software, such as the standalone desktop software Haploview (Barrett, Fry, Maller, & Daly, 2005), or for focused regions using the web-based application LocusZoom (Pruim et al., 2010). Conversely, `qqman` is distributed as an R package with no other dependencies that can be easily integrated into existing R-based scripted workflows to further enable automated reproducible research. Furthermore, users can take advantage of R’s very granular control of graphical output, enabling a high degree of customizability in creating high-resolution, publication-ready figures.

## ACKNOWLEDGEMENTS

The author wishes to thank all the hundreds of blog and Twitter commenters for pointing out bugs and other issues. The author extends a special thanks to Dan Capurso and Tim Knutsen for several useful code contributions and bug fixes.

